# Computational modeling reveals a hydrophobic force-induced pore-forming mechanism for cholesterol-dependent cytolysins

**DOI:** 10.1101/754705

**Authors:** Shichao Pang, Junchen Yang, Jingfang Wang

## Abstract

During the pore-forming process, cholesterol-dependent cytolysins (CDCs) bind to cholesterol-rich membranes and subsequently undergo a series of conformational changes, predominantly involving in the collapse of the protein and the transformation from α helices to β-hairpins to form a large hydrophobic pore. In the current study, we reconstructed a structural model for both the prepore and pore-forming complexes of PFO based on the cryo-EM data of pneumolysin and performed molecular dynamics simulations and free energy calculations to study the conformational changes in the PFO prepore-to-pore conversion. Our simulations indicate that D2 cannot collapse spontaneously due to the hydrogen bonding and pi-pi interactions between domains D2 and D3, which are partially weakened by binding to cell membranes and oligomerization. The free energy landscape for the prepore-to-pore conversion reveals that an additional force is required for the collapse of D2 to overcome an energy barrier of ∼ 24 kcal/mol. Based on these computational results, we proposed a hydrophobic force-induced pore-forming mechanism to explain the pore-forming process of CDCs. In this mechanism, the hydrophobic interactions between the TMHs and membranes are essential for the prepore-to-pore conversion. The hydrophobic force generated by the TMHs-membrane interactions drives the conformational changes in domains D2 and D3. These findings well explain how the conformational changes within two distant domains synergistically occur, and fits well for the previous biophysical and biochemical data.

## INTRODUCTION

Cholesterol-dependent cytolysins (CDCs) are a large family of bacterial pore-forming toxins that specifically bind to cholesterol in the plasma membrane of mammalian cells ^1, 2^. CDCs are usually expressed and secreted by Gram-positive bacterial pathogens as water-soluble monomer proteins with a molecular mass of 50-70 kDa ^3^. After binding to cholesterol, CDCs oligomerize and insert into the cell membrane with high concentrations of cholesterol (usually more than 25%) to form a large β-barrel pore with a diameter of 200-400 Å ^4^. The presence of cholesterol in the cell membrane is required for the pore-forming process of most CDCs. Because CDCs use cholesterol as their targets at the membrane surface ^5-7^. Besides, CDCs also display a high degree of sequence similarity, ranging from 40% to 80%, indicating that most CDCs exhibit a similar protein structure and thus share a common pore-forming mechanism. Thus, CDCs are essential for virus fusion, budding, and eukaryotic toxin mechanism ^8^.

Perfringolysin O (PFO) is secreted by the anaerobic bacteria Clostridium perfringens 9. This protein has been selected as a model system to study the CDC conformational alterations during their transition from soluble monomers to membrane-embedded pore complexes. The crystal structure of PFO is a soluble monomer composed of four domains, named domain 1 to domain 4 respectively (**Figure 1**) ^10^. Domain 1 (D1) shows a seven-stranded β-sheet structure with circumjacent α-helices. Domain 2 (D2) is composed of three long β-strands, linking domains 1 and 4. Domain 3 (D3) is an α/β/α-layered bundle, carrying two transmembrane β-hairpins (TMHs). Domain 4 (D4) is a compact β-sandwich, containing a membrane-sensing tryptophan-rich motif and a cholesterol recognition motif ^11^. The tryptophan-rich motif is an 11-residue peptide (also known as the undecapeptide or UDP), located in the C-terminal region of D4. This undecapeptide (ECTGLAWEWWR in PFO protein sequence) is highly conserved among CDC members, playing a crucial role in the CDC pore-forming process ^12, 13^.

**Figure 1.**
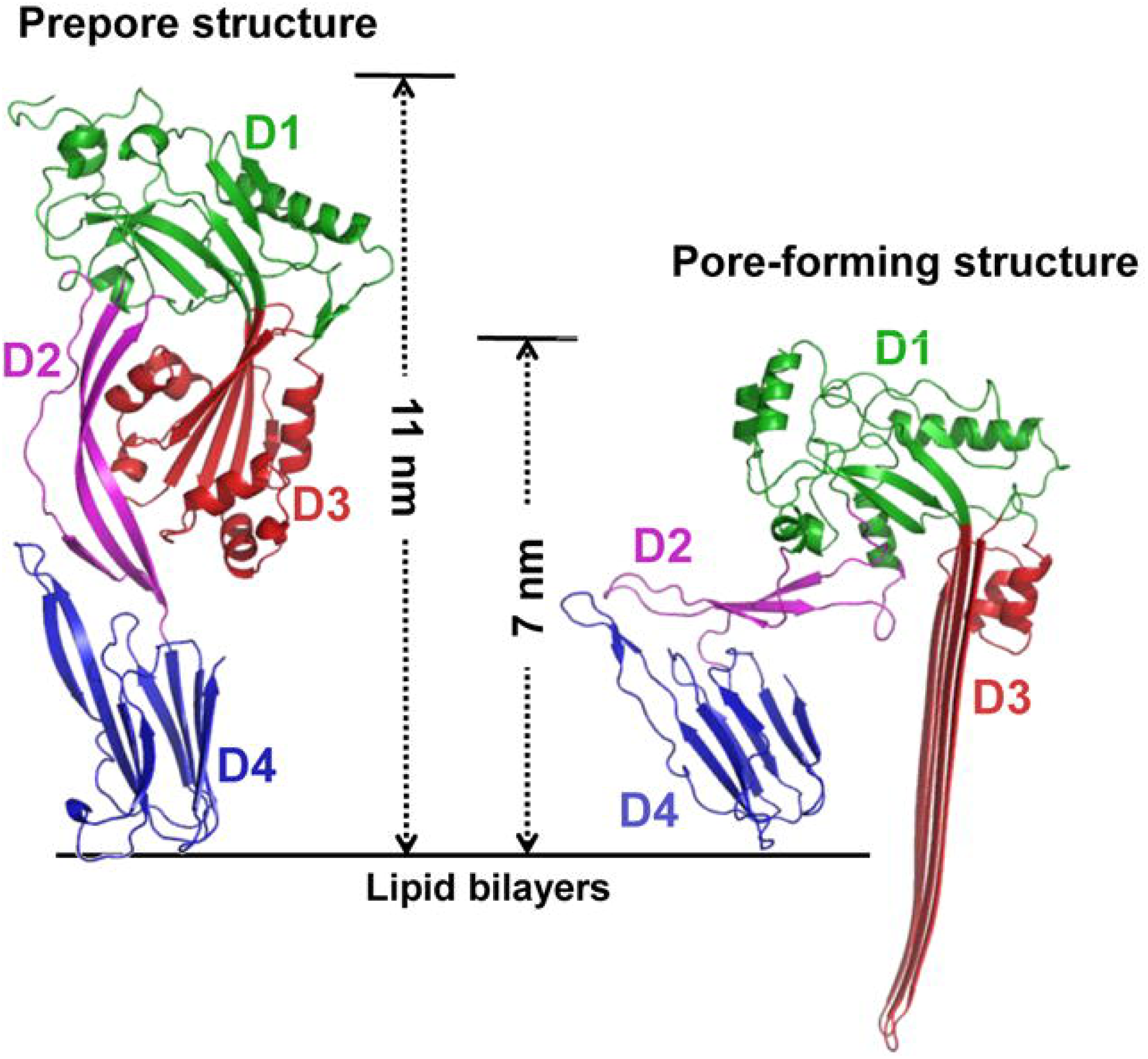
Prepore and pore-forming structural models of PFO. The prepore and pore-forming structural models were constructed based on the cryo-EM data of Pneumolysin prepore (EMD-1106) and pore complexes (EMD-4118). The PFO proteins are shown in cartoon representations with the structural domains colored in green, magenta, red, and blue, respectively. A significant distinction is detected in the protein height between the prepore and pore-forming structures, indicating a remarkable collapse in D2 during the pore-forming process.

The PFO pore-forming process contains two significant steps. The first step is the assembly of the prepore complex. In this step, the solution structure binds to the cell membrane via the tryptophan-rich motif and the cholesterol recognition motif, leading to a structural movement of D4 around a central β-sheet of D2 ^14-16^. The second step is the formation of the pore complex. In this step, a nearly 4 nm reduction in the protein height is detected due to the vertical collapse of D2. This observation is further confirmed by atomic force microscopy (AFM) ^17^, fluorescence-based distance measurements ^18^, and cryo-electron microscopy (cryo-EM)-based 3D reconstructions of *Streptococcus pneumoniae* pneumolysin (PLY) ^19^. Furthermore, the transformation from α-helices to β-hairpins occurs in the TMHs, assembling into a large β-barrel architecture and inserting into the lipid bilayers to form the pore, which is supported by fluorescence spectroscopy studies ^19, 20^.

Many biochemical and biophysical studies have been made to understand the large-scale structural changes in D2 and D3 ^17, 19-22^. However, it is still poorly understood what drives the vertical collapse of D2 and the β-barrel formation in D3 and how these structural alterations in D2 and D3 are coordinated. To answer these questions, we reconstructed the 3D structural models of the prepore and pore-forming structures for PFO based on the cryo-EM maps of the pneumolysin prepore and pore complexes, and employed long time-scale molecular dynamics (MD) simulations and free energy estimations to study the structural transitions during the prepore-to-pore conversion. We hope that our computational studies could be of general relevance to the pore-forming complexes of CDCs, and provide useful information for understanding the bacterial pore-forming process.

## MATERIALS AND COMPUTATIONAL METHODS

### 1. Reconstruction of the PFO prepore structural model

The PFO prepore structure used in the current study was reconstructed based on the crystal structure of PFO solution monomer (PDB accession no: 1pfo) ^10^ and cryo-EM map of the pneumolysin prepore complex (PDB accession no: EMD-1106) ^19^. Initially, each PFO domain in the solution structure was fitted into the cryo-EM map of the pneumolysin prepore complex as a rigid body using COOT ^23^. The flexible loops and unfolded regions were subsequently added manually, followed by geometry regularization in COOT. Finally, the initial model for the prepore structure was subjected to energy minimization by steepest descent for 5000 steps with the protein backbone fixed, followed by conjugate gradient for the next 5000 steps without any restriction.

### 2. Reconstruction of the PFO pore-forming structural model

Similar to the prepore structure, the pore-forming structure was also constructed based on the crystal structure of PFO solution monomer (PDB accession no: 1pfo) ^10^ and cryo-EM structure of the pneumolysin pore complex (PDB accession no: EMD-4118) ^24^. Differentially, D1, D2, D3 without the TMHs, and D4 in the solution structure were fitted into the cryo-EM map as rigid bodies using COOT. The TMHs were then modeled as β-hairpins and fitted into the cryo-EM map of the pneumolysin pore complex. The flexible regions were added manually, followed by geometry regularization in COOT. Finally, the structural model for the pore-forming structure was subjected to energy minimization by steepest descent for 5000 steps with protein backbone atom fixed, followed by conjugate gradient for the next 5000 steps without any restriction.

### 3. Reconstruction of the PFO prepore complex

Based on the prepore structure mentioned above, 36 PFO monomers were assembled and fitted into the cryo-EM map of the pneumolysin prepore complex (PDB accession no: EMD-1106) in UCSF Chimera ^25^, to form the prepore complex. The prepore complex was subsequently subjected to energy minimization by steepest descent for 5000 steps with protein backbone atom fixed, followed by conjugate gradient for the next 5000 steps without any restriction. The monomer-monomer interactions were finally abstracted from the adjacent monomers in the prepore complex.

### 4. Molecular dynamics simulation protocols

All non-polar hydrogen atoms in computational models were removed. Residue-specific pKa values were calculated by the linear Poisson-Boltzmann equation with a dielectric constant of 4.0 26. Hydrogen atoms were re-added to protein structures based on the computational pKa values mentioned above. After solvated in the explicit TIP3P water surroundings, the computational models were subjected to energy minimization by steepest descent for 5000 steps, followed by conjugate gradient for the next 5000 steps. MD simulations were finally performed with periodic boundary conditions and NPT ensembles (310K and 1 bar) by NAMD 2.1.2 ^27^ and CHARMM force field parameters ^28^. Langevin thermostat and isotropic scaling were used to control the temperature and the pressure in the simulation systems with a time constant of 0.1 ps and 1.0 ps respectively ^29^. For solvent simulation, the isothermal compressibility was set to 4.5×10^-5^ per bar. To restrict chemical bonds, the SHAKE algorithm was applied with a tolerance of 10^-6 30^. The particle mesh Ewald (PME) algorithm was used to treat electrostatic interactions with an interpolation order of 4 and a grid spacing of 0.12 nm ^31^. The van der Waals interactions were treated with a 12 Å cut-off.

### 5. Steered molecular dynamics simulation protocol

Initially, the short loop regions (L2 and L3) of D4 were fixed to simulate the lipid-binding state of the proteins. Subsequently, a constant force of 100 cal/mol/Å^2^ was applied to the D3 mass center, which is equal to be almost 60 pN exerted on the 4-nm lipid bilayers. The direction of the applied force was selected as the paths from the D3 mass center to the membrane surface with a tilt of ∼ 15° from the long axis of the protein, to avoid D3 directly interacting with D4 during the simulation process. The steered MD simulation was finally performed by the module implemented in NAMD 2.1.2 with an integration step of 2 fs, and the coordinates were saved every 1 ps. After nearly 600-ps simulations, the prepore structure converted to the pore-forming structure with the collapse of D2.

### 6. Free energy calculations for the prepore-to-pore conversion

The energy landscape for the prepore-to-pore conversion was described by the potential of mean force (PMF) by the umbrella sampling method ^32^. In the prepore-to-pore conversion, a total of 50 windows were applied for the potential of mean force calculations. For each window, 50 snapshots were extracted from the simulation trajectory of the steered MD simulation mentioned above, acting as the initial structures for free energy calculations. These initial structures were subsequently subjected to a short MD simulation (nearly 2 ns) with the simulation protocols mentioned above, and the potential of mean forces was simultaneously calculated by using the PMF module implemented in NAMD 2.1.2 with CHARMM force field parameters.

### 7. Data analysis and structural visualization

The height of D2 was defined as the Cα distance of Gly72 and Ser88 in the protein structure. The bending angle of D2 was defined as the Cα angle of Gly72, Lys82, and Ser88. The height of D3 was measured as the distance between the D3 mass center and the membrane surface. The rotating angle of the D3 core β-strand structure was measured as the Cα angle of Lys272, Phe230, and Ser234. In the hydrogen bonds, the distance between the heavy atoms of hydrogen donors and hydrogen acceptors should be less than 3.5 Å ^33^. All the protein structures were visualized by PyMol ^34^.

The lifetime for the energy barrier was calculated by the following equation: *k*_0_ =A×e^((-ΔG)/*k*T), where A is the attempt frequency, ΔG is the energy barrier, *k* is Boltzmann’s constant and T is the temperature. According to the force spectroscopy studies ^35^, the attempt frequency A was set to 1×10^7^ per second. The energy barrier ΔG was set to 16.32 kcal/mol based on the free energy landscape of the prepore-to-pore conversion. Thus, the equation mentioned above yields a *k*_0_ of 6.9×10^-6^ per second, corresponding to a lifetime of nearly 1.5 days.

## RESULTS

### 1. D2-D3 interactions lock the TMHs in the prepore structure

Atomic force microscopy ^17^, förster resonance energy transfer (FRET) studies ^18^, and single-particle cryo-EM based 3D reconstruction of PLY ^19^ revealed a remarkable 4 nm reduction in the protein height during the prepore-to-pore conversion of PFO (**Figure 1**). Crystallographic studies indicated that the flexibility of D2 had a significant impact on the protein height of CDCs ^10, 15^. This domain is composed of three long β-strands, showing a high degree of flexibility than the other domains. The vertical collapse of the β-strand structures in D2 can cause a structural reduction in the protein height for CDCs. To study the structural flexibility of D2, we performed long-timescale MD simulations on the PFO prepore structure. As a result, no significant fluctuations are detected in the height and the bending angle of D2 during our simulations (Supplementary **Figure S1** and **Figure S2**). This is mainly because of the significant hydrogen-bond network between the backbone residues of D2 and the TMHs in D3 (**Figure 2A-2C**). This hydrogen-bond network not only locks the TMHs in D3 to avoid the transformation from α-helices to β-hairpins but also prevents the collapse of D2.

**Figure 2.**
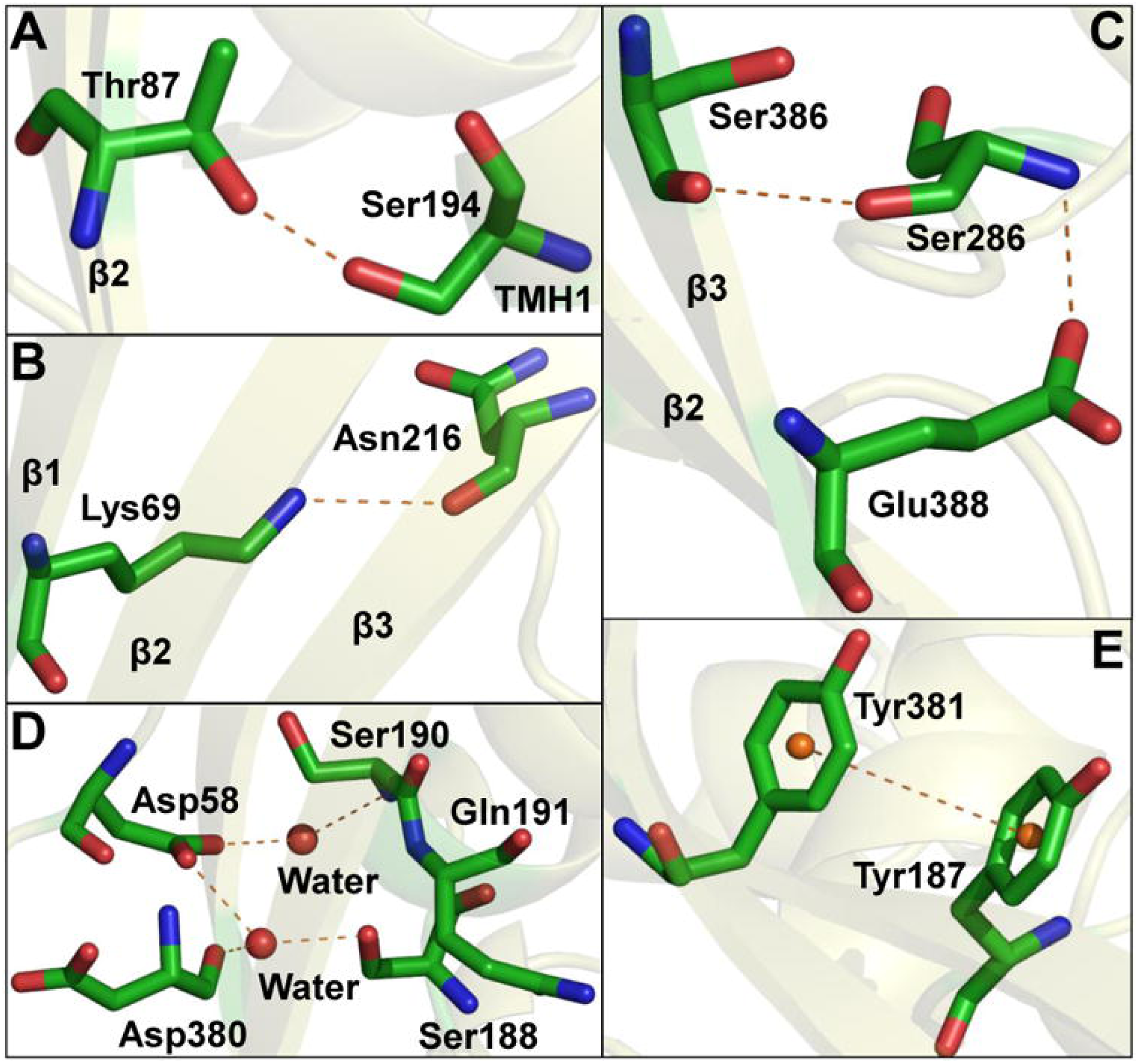
A stereoview of the D2-TMHs interactions. (A) Ser194 in TMH1 interacts with the backbone residue Thr87 in D2 via a hydrogen bond with an occupancy of 17.2%. (B) Asn216 in D3 and Lys69 in D2 forms a hydrogen bond with an occupancy of 27.5%. (C) Ser286 forms hydrogen bonds with the backbone residues Ser386 and Glu388 in D2 (with occupancies of 20.2% and 25.9%, respectively). (D) Water-mediated hydrogen bonds among Asp58, Ser188, Ser190, and Asp380. (E) Pi-pi stacking interaction between the aromatic sidechains of Tyr187 and Tyr381.

Compared with the solution structure, the hydrogen-bond network in the prepore structure is slacked down (**Table 1**). Except for the one formed by Lys69 and Asn216, the occupancies for all the other hydrogen bonds are decreased to below 10%, partly increasing the flexibility of the TMHs. Asn216 is located in the loop region between TMH1 and β-strands in D3, and its hydrogen bonding with Lys69 functions to stabilize the flexible loop region in D3. Mutagenesis experiments and AFM imaging studies show that the direct contacts between the backbone residues in D2 and D3 are adverse to the pore-forming transition ^21, 36^. Structural analysis indicates that the loss of contacts between D2 and D4 in the prepore structure results in the lessened hydrogen-bond network. X-ray crystallography and computational studies of different conformational states of PFO suggest that membrane binding leads to a significant structural movement between D2 and D4 ^15, 22^. Compared to the soluble monomer, D4 twists nearly 15° in the prepore structure, resulting in the loss of contacts between D2 and D4. This finding also fits well with the crystal structure analysis, cryo-EM based 3D reconstruction, and computational simulations ^15, 19, 22^.

**Table 1.** The occupancies of the hydrogen bonds between the backbone residues in D2 and D3.

| No. | H-Donor | H-Acceptor | Occupancy |  | p-value |
| --- | --- | --- | --- | --- | --- |
|  |  |  | Solution | Prepore |  |
| 1 | Ser194 | Thr87 | 10.2% | 3.6% | 0.007 |
| 2 | Lys69 | Asn216 | 15.5% | 10.3% | 0.010 |
| 3 | Ser386 | Ser286 | 15.2% | 4.5% | 0.005 |
| 4 | Ser286 | Glu388 | 12.9% | 4.1% | 0.009 |

In addition to the hydrogen-bond network formed by the backbone residues, water-mediated hydrogen bonds and pi-pi stacking formed by the aromatic residues also contribute to the D2-D3 interactions. Significant water-mediated hydrogen bonds are detected around Asp58, Ser188, Ser190, and Asp380 (**Figure 2D**). A water molecule links Asp380 with Asp58 and Ser188 respectively, and another water molecule links Asp58 with Ser190. A significant pi-pi stacking is detected between the aromatic sidechains of Tyr187 and Tyr381 (**Figure 2E**), functioning to position the large sidechains of the backbone residues.

### 2. PFO oligomerization releases the TMHs from D2

Many biophysical studies indicate that unlocking the TMHs from the β-strands in D2 is the primary requirement for the prepore-to-pore conversion of CDCs. To release the TMHs, PFO needs to form a stable oligomerized complex, breaking the hydrogen-bond network and pi-pi stacking interactions between D2 and D3 ^37^. In the PFO dimer, the monomer-monomer interactions make the core β-strand structures of D3 twist approximately 15°. To support this point, we defined the rotating angle of the core β-strand structure in D3 as the Cα angle of Lys272, Phe230, and Ser234. As a result, the rotating angle in the PFO monomer is 95.0 ± 8.5°, statistical-significantly different from the dimer (110.4 ± 6.5°, t-test p-value < 0.05, as shown in Supplementary **Figure S3**). The increased rotating angle in the PFO dimer releases the TMHs from the D2-D3 interactions. However, the core β-strands of D3 are titled by 20° in the pore-forming structure, in good agreement with the observations in suilysin and pneumolysin pore complexes ^24, 38^.

The release of the TMHs in D3 also plays a crucial role in stabilizing the PFO dimer and in the transformation from α-helices to β-hairpins in the prepore-to-pore conversion ^14^. This α-helices to β-hairpins transformation begins with the structural rearrangement of β5 in D3, which is located in the core β-strand structure of D3 adjacent to TMH2 (**Figure S4**) ^37^. Because Lys336 in β5 forms an intermolecular electrostatic interaction with Glu183 in β1 in the adjacent monomer (**Figure 3A** and **Figure 3B**). In the pore-forming structure, Glu183 is located in β1 in the TMH1. Thus, the prepore-to-pore conversion requires the disruption of the electrostatic interaction formed by Glu183 and Lys336, to release the TMHs in the PFO dimer. The flexible helices (TMH1 and TMH2) make β5 depart from β4 and expose the edge of β4. As a result, the backbone residues of β4 and β1 in the adjacent monomer form multiple hydrogen bonds.

**Figure 3.**
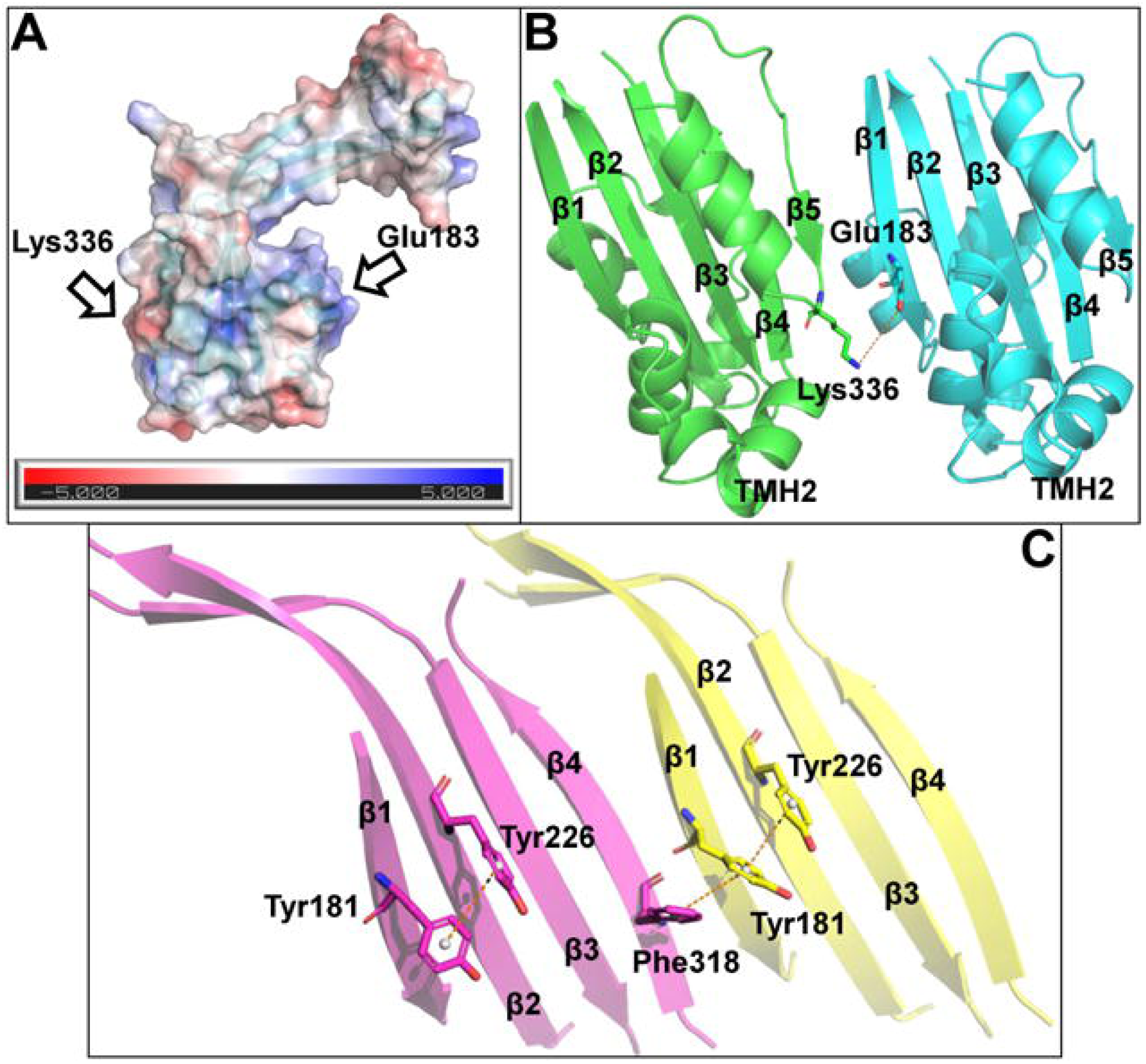
Monomer-monomer interactions in the D3 domain. (A) Electrostatic map of the D3 domain. The charged residues Glu183 and Lys336 are labeled in the electrostatic map. (B) The stereo-view of the Glu183-Lys336 interaction in the prepore complex. (C) Pi-pi stacking interactions in the transmembrane β-hairpins in the pore-forming complex.

Furthermore, the aromatic side-chains of Tyr181 in β4 and Phe317 in β1 in the adjacent monomer also form an intermolecular pi-pi-stacking interaction (**Figure 3C**). Both hydrogen bonding and pi-pi stacking interactions stabilize the rearranged core β-strand structure in D3, having significant impacts on the formation of the transmembrane β-hairpins.

### 3. Free energy landscape for the prepore-to-pore conversion

After releasing the TMHs, PFO undergoes a prepore-to-pore conversion with the collapse of D2 and the transformation from α-helices to β-hairpins in the TMHs. To study these structural changes, we used steered MD simulations and the umbrella sampling method to describe the free energy landscape of the D2 collapse (**Figure 4**). In the free energy landscape, a global minimum and a local minimum are located at the reaction coordinate distance of 1.5 Å and 45 Å, respectively. The former is associated with the prepore structure, and the latter is presumed to associate with the pore-forming structure. An energy barrier of 24.34 kcal/mol (centered at ∼ 29 Å) appears between the global and local minimums. This observation is in good agreement with the AFM imaging studies ^17, 36^, indicating that an additional force is required to reduce the energy barrier for the prepore-to-pore conversion.

**Figure 4.**
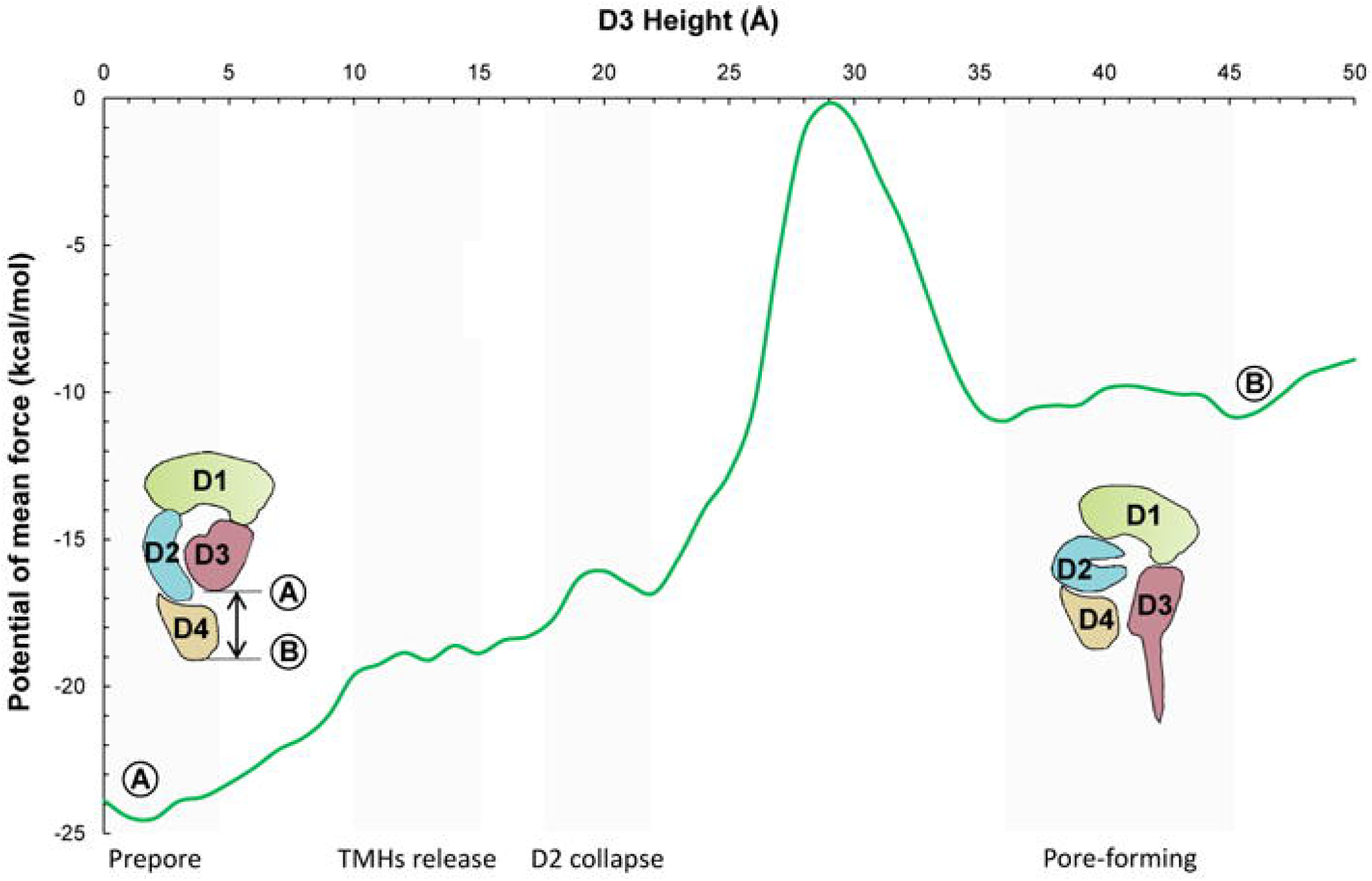
Free energy landscape for the prepore-to-pore conversion. The global minimum at ∼ 1 Å is associated with the prepore structure, and the local minimum at the height of ∼ 45 Å is associated with the pore-forming structure. An energy barrier of 24.34 kcal/mol separates the global and local minimums.

Notably, there is a small peak at the reaction coordinate distance of 17-22 Å with an energy barrier of 16.32 kcal/mol in the energy landscape. This energy barrier is relevant to the collapse of D2. Both the reaction coordinate distance (about 5 Å) and the energy barrier (16.32 kcal/mol) for this peak well fit with the AFM measurement and mutagenesis experiments ^36^. According to the force spectroscopy studies ^35^, the attempt frequency is about 10^7^ per second. Thus, the energy barrier for the collapse of D2 (16.32 kcal/mol) yields to a lifetime of nearly 1.5 days (for detailed information, please see Materials and Computational Methods). These findings reveal that the energy barrier for the collapse of D2 cannot be overcome by the thermal dynamics alone.

The peaks at the reaction coordinate distance of 10-15 Å is presumed to be associated with the release of the TMHs. As mentioned above, the release of the TMHs requires the disruption of D2-D3 interactions. The energy barrier between the global minimum and these peaks is nearly 6 kcal/mol, which is used for breaking the hydrogen-bond and pi-pi interactions between domains D2 and D3.

Near the local minimum, there are some small rises in the energy at the reaction coordinate distance of 40-45 Å. This energy barrier is associated with the D2-D4 interactions in the pore-forming structure. The collapsed D2 domain in the pore-forming structure employs more direct interactions with D4 than the prepore structure. For instance, Lys69 and Asn73 in the loop region of D2 form a hydrogen bond with Tyr415 and Asn488 in D4, respectively (**Figure 5**). Of note, Ser386 and Glu388 in D2 simultaneously form a hydrogen bond with Asn480 in the loop region of D4 in the pore-forming structure (**Figure 5**). However, these residues function to lock the TMH2 of D3 in the prepore structure.

**Figure 5.**
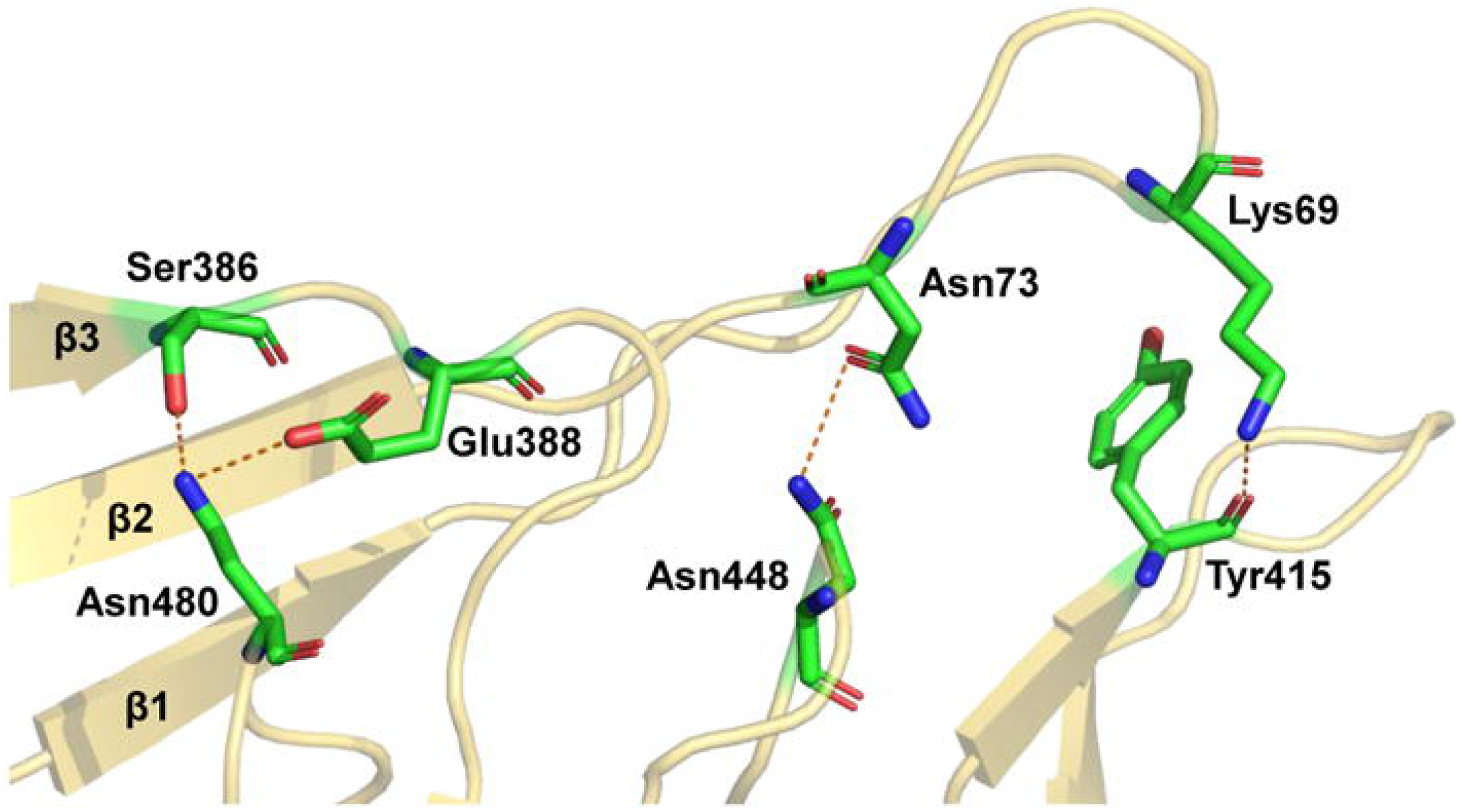
The stereo-view of the hydrogen-bond network in the D2-D4 interface. The collapsed D2 directly interacts with the flexible loop region of D4, forming four significant hydrogen bonds. These hydrogen bonds stabilize the β-strands and the flexible loop regions in D2 and D4.

## DISCUSSION

The prepore-to-pore conversion of CDCs requires the collapse of D2. However, the free energy landscape for the prepore-to-pore conversion indicates that the collapse of D2 cannot spontaneously happen. What drives the vertical collapse of D2 and the β-barrel formation in D3? As transmembrane helices, TMH1 and TMH2 have many hydrophobic residues exposed in the solvent (**Figure 6**). As known, the hydrophobic groups employ specific interactions in the water surrounding, which can be described by the solvation energy. According to the previous theoretical studies and Monte Carlo simulations, the solvation energy for the hydrophobic peptides is about −96 ± 6 cal/mol/ Å^2^, which is equal to 1.43 pN/Å^2 39^. The lipid bilayers are almost 4 nm thick ^40^, thus the hydrophobic force between the TMHs and lipid bilayers is about 57 pN. In the human body, the forces generated by the myosin are 5-6 pN ^41^, while the forces for RNA polymerases and the portal motos are nearly 25 pN and 50 pN, respectively ^42, 43^. The hydrophobic force generated by the TMHs-lipid bilayers interactions is almost the same as the portal motor, indicating that it is strong enough to induce the collapse of D2 and the transformation from α-helices to β-hairpins in D3.

**Figure 6.**
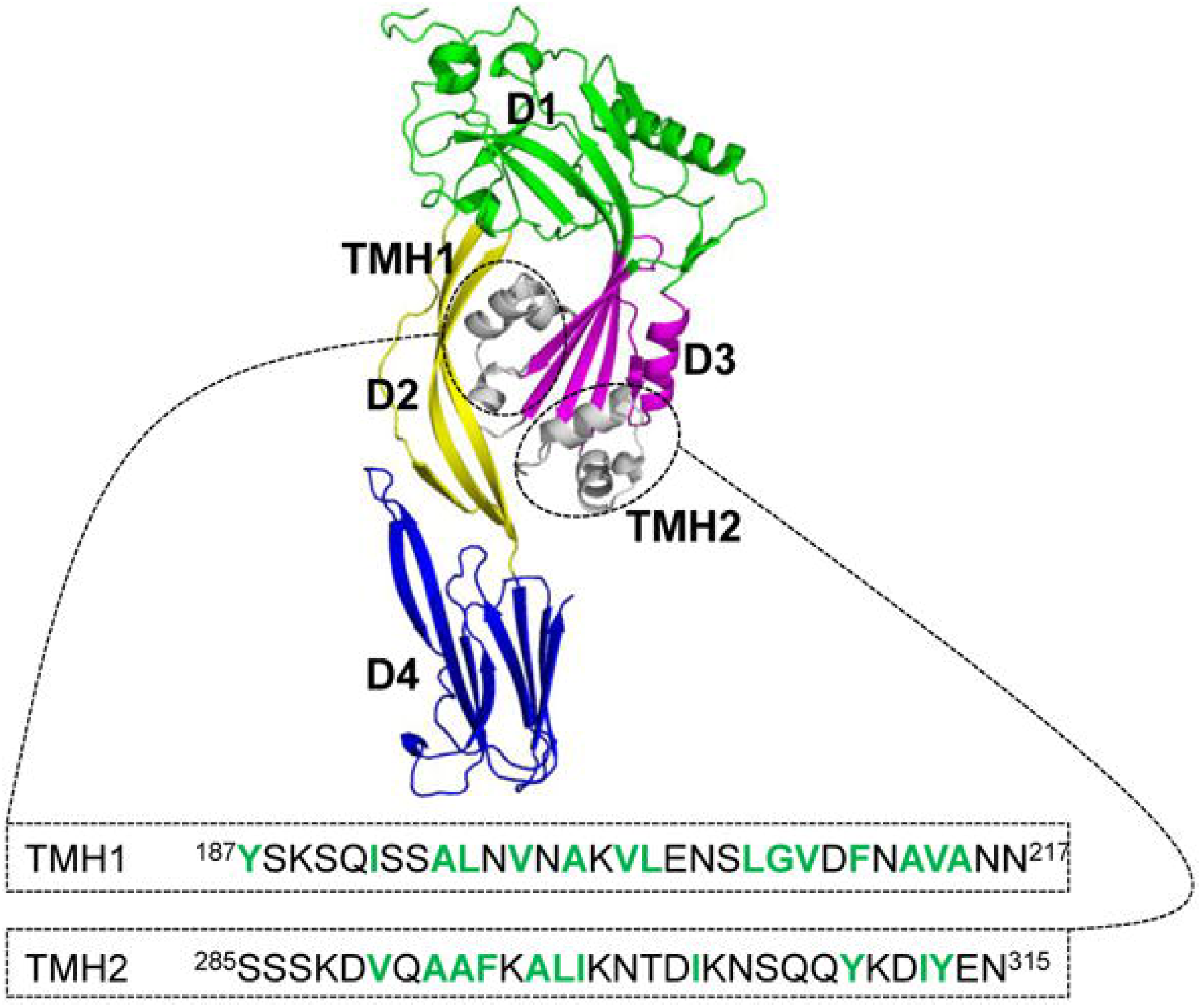
The amino acid composition of the TMHs. As transmembrane helices, TMHs contains a large number of hydrophobic residues, whose solvation energy is about −96 ± 6 cal/mol/ Å^2^, equal to 1.43 pN/Å^2^. Thus, the TMHs-lipid bilayers will generate an additional force of about 57 pN, almost the same as the portal motor.

## CONCLUSION

The pore-forming process of CDCs usually contains four steps: binding to membranes and assembly of the prepore complex, dimerization, the vertical collapse of D2, and the transformation from α-helices to β-hairpins in D3. Although many good attempts have been made to study these steps, it is still unknown what drives the collapse of D2 and the structural transformation in the TMHs. In the current study, we proposed a hydrophobic force-induced pore-forming mechanism to explain the collapse of D2 and the transformation from α-helices to β-hairpins in D3. Due to a large number of hydrophobic residues in the TMHs, the TMHs-lipid bilayer interactions generate a hydrophobic force of about 57 pN, which is strong enough to induce the collapse of D2 and the transformation from α-helices to β-hairpins in D3. We hope that these findings may be of general relevance to the bacterial pore-forming complexes, providing useful information for engineering specifically tailored pores for biotechnological applications.

## FIGURE CAPTION

**Figure S1.** The flexibility of D2 in the prepore structure. RMS deviations for the protein backbone are used to assess the flexibility of D2. In the prepore structure, the RMS deviations for the backbone structure is 4.64 ± 0.44 Å with a maximum fluctuation of 2.19 Å.

**Figure S2.** The height and bending angle of D2 in the prepore structure. The height of D2 is defined as the Cα distance of Gly72 and Ser88 in the PFO protein structure. The bending angle of D2 is defined as the Cα angles of Gly72, Lys82, and Ser88. The height and bending angle of D2 in the prepore structure are 42.8 ± 1.2 Å and 123.5 ± 4.4° along with MD simulations.

**Figure S3.** Rotating angles of the D3 core β-strand structure in the monomer (red curve) and dimer (blue curve). The rotating angle is defined as the Cα angle of Lys272, Phe230, and Ser234 in the D3 core β-strand structure. In the PFO dimer, the rotating angle of the D3 core β-strand structure is 110.4 ± 6.5°, nearly 15° larger than the monomer (95.0 ± 8.5°).

**Figure S4**. The stereo-view of the D3 core β-strand structure. The D3 core structure is composed of four β-strands contiguous with two α-helical bundles that ultimately form the TMHs. This formation is induced by the disengagement of β5 from β4 to expose the edge of β4 in one monomer, resulting in multiple backbone hydrogen bonds with β1 of the adjacent monomer and an intermolecular pi-pi stacking interaction with Tyr181 in one monomer and Phe318 in the adjacent monomer.

## ASSOCIATED CONTENT

### Supporting Information

The following files are available free of charge.

**Figure S1.** The flexibility of D2 in the prepore structure. RMS deviations for the protein backbone are used to assess the flexibility of D2. In the prepore structure, the RMS deviations for the backbone structure is 4.64 ± 0.44 Å with a maximum fluctuation of 2.19 Å. (file type, PDF)

**Figure S2.** The height and bending angle of D2 in the prepore structure. The height of D2 is defined as the Cα distance of Gly72 and Ser88 in the PFO protein structure. The bending angle of D2 is defined as the Cα angles of Gly72, Lys82, and Ser88. The height and bending angle of D2 in the prepore structure are 42.8 ± 1.2 Å and 123.5 ± 4.4° along with MD simulations. (file type, PDF)

**Figure S3.** Rotating angles of the D3 core β-strand structure in the monomer (red curve) and dimer (blue curve). The rotating angle is defined as the Cα angle of Lys272, Phe230, and Ser234 in the D3 core β-strand structure. In the PFO dimer, the rotating angle of the D3 core β-strand structure is 110.4 ± 6.5°, nearly 15° larger than the monomer (95.0 ± 8.5°). (file type, PDF)

**Figure S4**. The stereo-view of the D3 core β-strand structure. The D3 core structure is composed of four β-strands contiguous with two α-helical bundles that ultimately form the TMHs. This formation is induced by the disengagement of β5 from β4 to expose the edge of β4 in one monomer, resulting in multiple backbone hydrogen bonds with β1 of the adjacent monomer and an intermolecular pi-pi stacking interaction with Tyr181 in one monomer and Phe318 in the adjacent monomer. (file type, PDF)

## ABBREVIATIONS

CDC: Cholesterol-dependent cytolysin,
PFO: Perfringolysin O,
UDP: Undecapeptide,
AFM: Atomic force microscopy,
Cryo-EM: Cryo-electron microscopy,
PLY: Pneumolysin,
FRET: Förster resonance energy transfer,
TMHs: Transmembrane β-hairpins,
MD: Molecular dynamics,
PME: Particle mesh Ewald,
PMF: Potential of mean force,

