## Supplementary figures and images for "Computational modeling reveals a hydrophobic force-induced pore-forming mechanism for cholesterol-dependent cytolysins"

### Figure S1

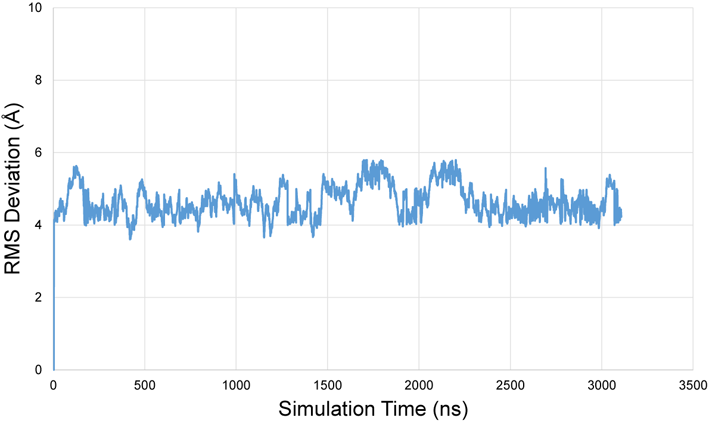

### Figure S2

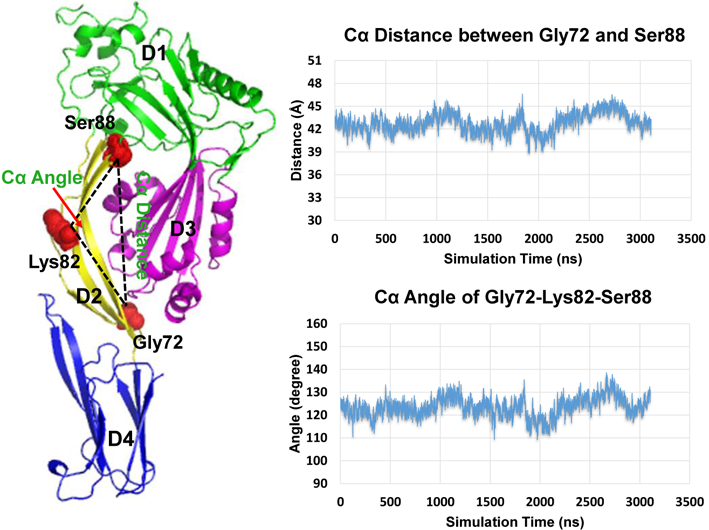

### Figure S3

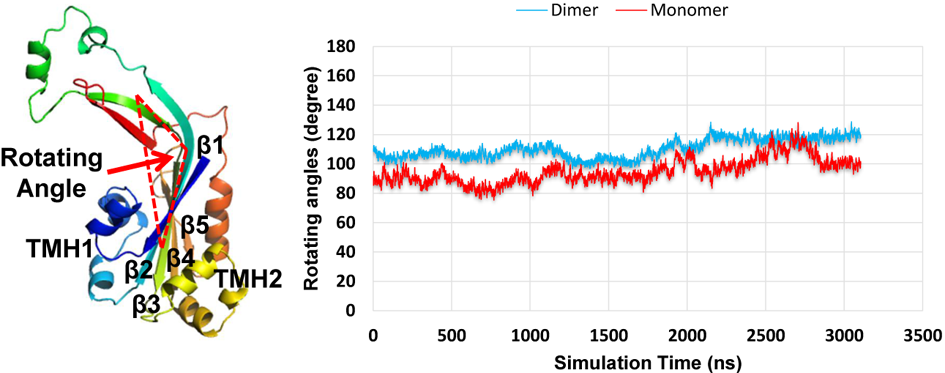

### Figure S4

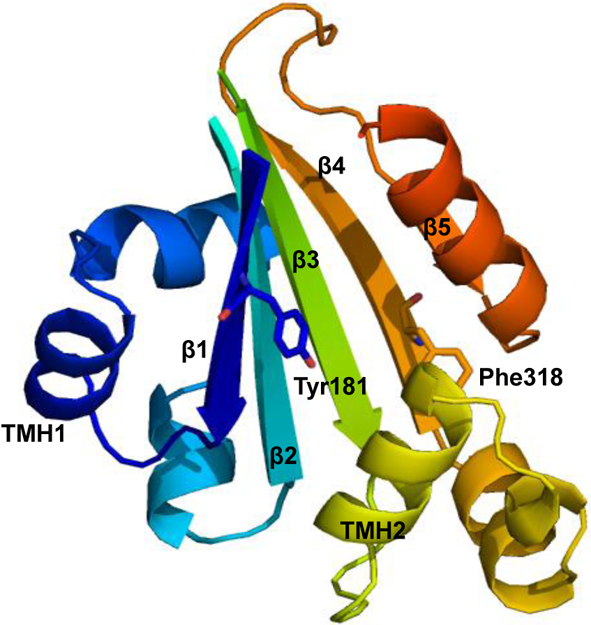
